# Estrus-Cycle Regulation of Cortical Inhibition

**DOI:** 10.1101/314641

**Authors:** Ann M. Clemens, Constanze Lenschow, Prateep Beed, Lanxiang Li, Rosanna Sammons, Robert K. Naumann, Hong Wang, Dietmar Schmitz, Michael Brecht

## Abstract

Female mammals experience cyclical changes in sexual receptivity known as the estrus-cycle. Little is known about how estrus affects the cortex although alterations in sensation, cognition and the cyclic occurrence of epilepsy suggest brain-wide processing changes. We performed *in vivo* juxtacellular and whole-cell recordings in somatosensory cortex of female rats and found that the estrus-cycle potently altered cortical inhibition. Fast-spiking interneurons strongly varied their activity with the estrus-cycle and estradiol in ovariectomized females, while regular-spiking excitatory neurons did not change. *In vivo* whole-cell recordings revealed a varying excitation-to-inhibition-ratio with estrus. *In situ* hybridization for estrogen receptor β (Esr2) showed co-localization with parvalbumin-positive interneurons in deep cortical layers, mirroring the laminar distribution of our physiological findings. *In vivo* and *in vitro* experiments confirmed that estrogen acts locally to increase fast-spiking interneuron excitability through an estrogen receptor β mechanism. We conclude that sex hormones powerfully modulate cortical inhibition in the female brain.

## Introduction

Female mammals experience cyclical changes in physiology and behavior, which are mediated by hormonal fluctuations targeting the reproductive organs and brain (Beach, 1976; Erskine, 1989; Fink et al., 2012; Pfaff, 1999). In humans, generalized changes in brain function include alterations of sensory processing (Penton-Voak and Perrett, 2000), cognitive function (Sundström Poromaa and Gingnell, 2014) and the periodic occurrence of epilepsy (Pennell, 2009) during the menstrual cycle. While cyclical changes have been studied in detail in subcortical structures (Blume et al., 2017; Micevych and Meisel, 2017), relatively little is known about how the estrus-cycle affects cortical processing. Since the pioneering work of Hubel and Wiesel revealed the remarkable stimulus selectivity in sensory cortical neurons (Hubel and Wiesel, 1977), investigations of the sensory cortices have shaped our thinking about information processing in the brain. A large body of work revealed that attention affects cortical discharges and has small effects (typically in the 10% range) in primary cortices and larger effects in higher order cortices (Maunsell and Cook, 2002; Yoshor et al., 2007). In our own work on social touch we observed an unexpected and large difference in socially evoked discharges between recordings in male and female animals (Bobrov et al., 2014). These observations and earlier work on estrus-related changes in spine turnover in hippocampal (Gould et al., 1990) and cortical circuits (Chen et al., 2009) suggested a role of sex hormones in shaping cortical processing and prompted us to examine the effects of the female estrus-cycle on cortical activity. Using awake *in vivo* juxtacellular and *in vivo* whole-cell patch clamp recordings in freely-cycling and ovariectomized female rats, we find dramatic and highly cell-specific effects of the female estrus-cycle and the sex hormone estrogen on cortical activity. *In situ* hybridization with parvalbumin (PV) co-labeling revealed co-localization of Esr2 and PV positive neurons in deep cortical layers which mirrored the depth distribution of our physiological data. Finally, we performed local application of 17 beta-estradiol to the cortex *in vivo* and *in vitro* and found that estrogen acts locally in the cortex to increase excitability in parvalbumin-positive fast-spiking neurons through an estrogen receptor β (Esr2) dependent mechanism.

## Results

We investigated the influence of sex and cell-type on the somatosensory cortex in a staged social facial touch paradigm where we presented stimulus rats to a head-fixed subject rat (Lenschow and Brecht, 2015). We performed juxtacellular recordings with cellular labeling while facial interactions where filmed from above and illuminated with an infrared light (**Figure 1A**). Fast-spiking and regular-spiking neurons were differentiated based on morphology and characteristics of spike shape (**Figure 1B**). All morphologically (non-pyramidal somato-dendritic architecture, smooth dendrites) identified interneurons fit the fast-spiking spike-shape criteria (4 out of 4; 100%) and a majority of morphologically (pyramidal somato-dendritic architecture, spiny dendrites) identified regular-spiking neurons were confirmed as principal cells (5 out of 6; 83%; **Figure 1C**). An identified fast-spiking interneuron with a non-pyramidal somato-dendritic morphology and strong responses to social touch is shown in **Figure 1D.** An identified regular-spiking principal cell with a pyramidal somato-dendritic morphology and weaker responses to social touch is shown in **Figure 1E.** When we examined the cell-specificity of responses to social facial touch, we observed on average greater responses in fast-spiking neurons compared with regular-spiking neurons. This was apparent in identified neurons (**Figure 1D–E**) as well as across the population of recorded neurons (**Figure 1F**). Fast-spiking neurons responded more to social touch in both male and female subject rats and responses did not depend on the sex of the stimulus rat (data not shown).

**Figure 1:**
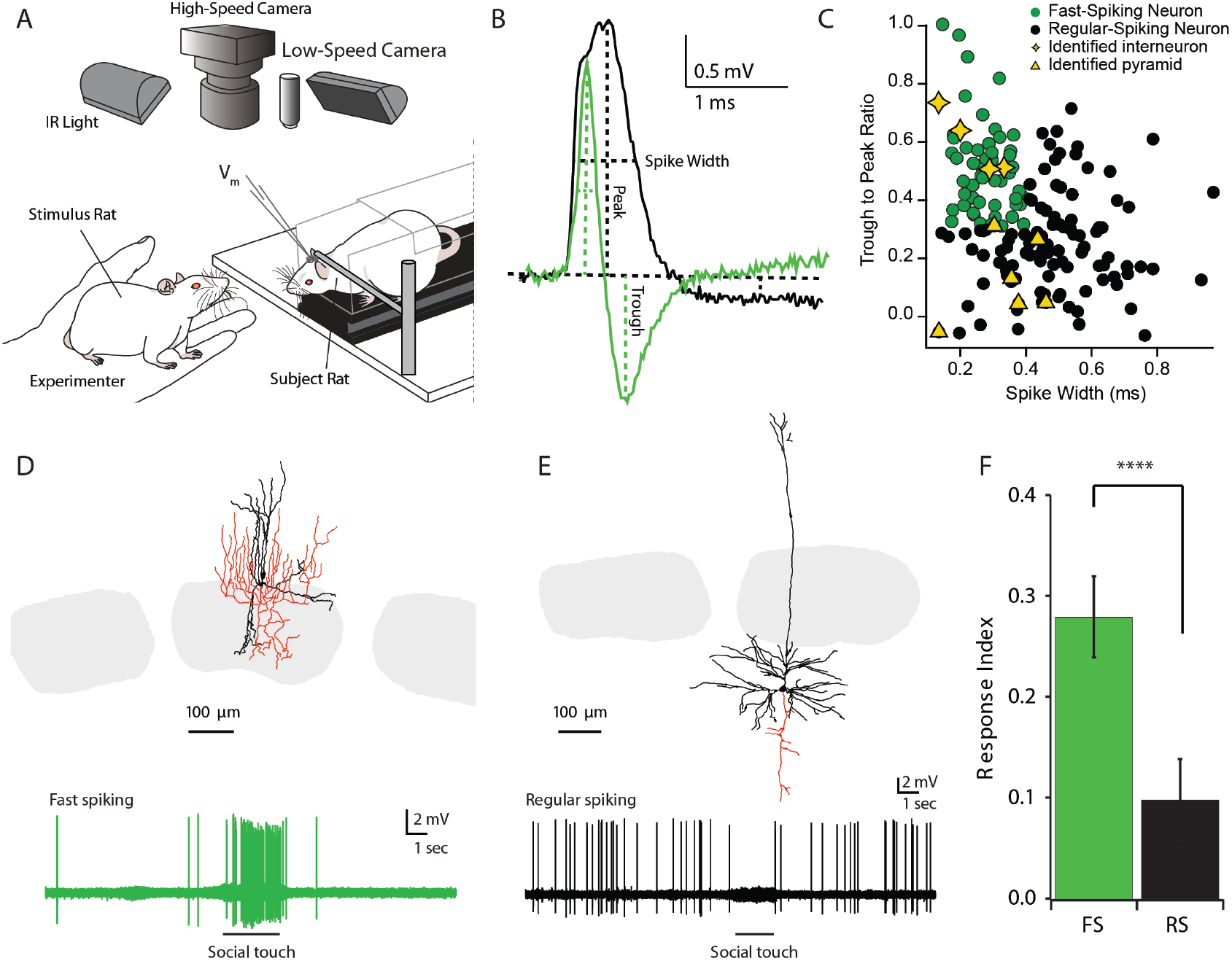
Juxtacellular identification of fast- and regular-spiking neurons and responsiveness to social facial touch in rat barrel cortex. **A**, Setup for awake juxtacellular recordings of social facial interactions with a head-fixed subject rat positioned below high-speed and low-speed cameras as well as an infrared light for illumination. Stimulus rats were presented by hand by the experimenter. **B**, Fast- and regular-spiking neurons were classified based on the width of the spike at half-maximal amplitude and the ratio of the amplitude of the spike trough to the amplitude of the spike peak. **C**, Cells were labeled and recovered when possible. Fast-spiking neurons (spike-width < 0.4 ms and trough-to-peak ratio > 0.3) were identified morphologically as interneurons in 4 out of 4 cases. Regular-spiking neurons (spike-width > 0.4 ms or trough-to-peak ratio < 0.3) were identified morphologically as pyramidal neurons in 5 out of 6 cases (FS: N=138 unique interactions; RS: 359 unique interactions). **D**, A morphologically confirmed fast-spiking neuron and the response of this cell to social facial touch. Dendritic processes are shown in black, axonal processes in red. Barrels are shown in gray. **E**, An identified regular-spiking pyramidal neuron and the response of this cell to social facial touch. Barrels are shown in gray. **F**, Fast-spiking (FS) neurons were overall more responsive to social facial touch when compared with regular-spiking (RS) neurons. Response indices indicate the normalized change from ongoing to the period during social facial touch (FS: N=177 interactions; RS: N=363 interactions; recordings from male and female subject animals; Mann-Whitney *U* test p<0.001). ****p<0.001.

Socio-sexual behavior in females is highly dependent on estrus state (Beach, 1976; Erskine, 1989; Young et al., 1941). We assessed whether the observed differential cortical responses to social interaction were dependent on the hormonal cycle. We used multiple methods to assess the estrus state of the females and found that a combination of measures was most reliable. At the same time daily, vaginal smears were collected and vaginal impedance was measured. We found that these two measures corresponded with each other (**Figure 2A**) and each period of the cycle spanned 3–8 days. The period of sexual receptivity in female rats begins during vaginal proestrus and continues into the day of vaginal estrus (Young et al., 1941) (indicated by pink, **Figure 2A**), the period known as “Estrus”, therefore, encompasses the periods of vaginal proestrus and vaginal estrus. Vaginal metestrus and diestrus were considered the “Non-estrus” period. The period of “Estrus” or sexual receptivity, marked by a rise in the sex hormone estrogen, is followed by a rise in progesterone^18^ (**Figure 2B**). To our surprise, when recordings were sorted relative to the estrus-cycle, we found that non-estrus fast-spiking neurons had lower ongoing firing rates (**Figure 2C**) as compared to fast-spiking neurons recording during estrus (**Figure 2D**). The dependence of ongoing firing on estrus day was observed in the average of all recorded females and showed a cyclic-relationship when plotted relative to the day of vaginal estrus (**Figure 2E**) as well as when cells were separated into estrus and non-estrus groups. The estrus changes in ongoing activity in fast-spiking neurons amounted to a dramatic three-fold increase in firing rate (**Figure 2F**). We did not find a significant change in responsiveness during the period of social touch between estrus and non-estrus states in fast-spiking neurons (**Supplementary Figure 1A–B**).

In regular-spiking neurons, ongoing firing (**Figure 2G–J**) and responsiveness (**Supplementary Figure 1C–D**) were low and showed no significant difference between “Estrus” and “Non-Estrus” periods.

**Figure 2:**
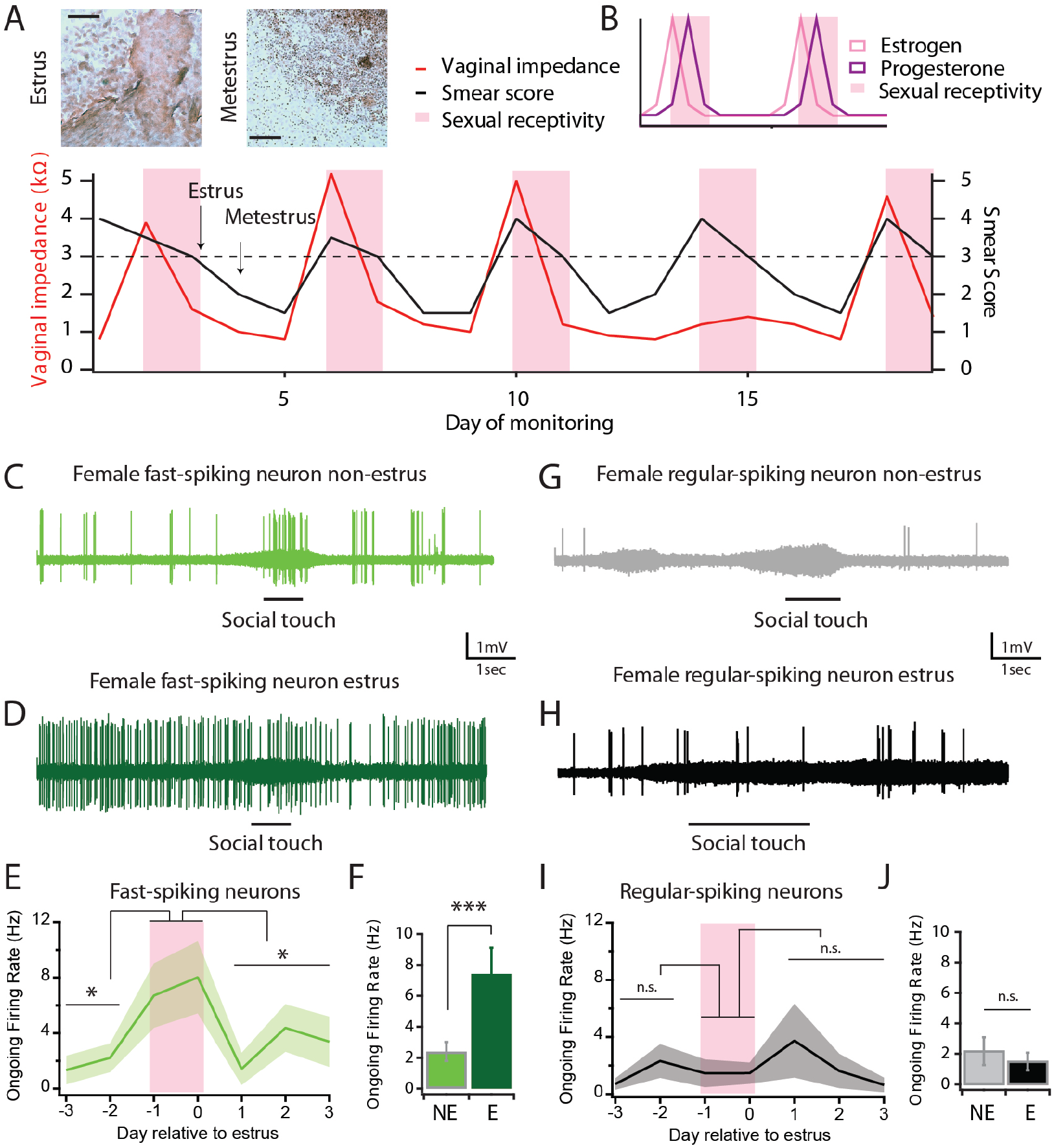
Activity of fast-spiking neurons is regulated by the estrus-cycle. **A**, Methodology used to assess the estrus state of female rats. Example images of estrus and metestrus vaginal smear (Scale bar: 200 μm). Estrus smears were scored based on cellular composition of vaginal smears (Smear Score: 1: Diestrus, 2: Metestrus, 3: Estrus, 4: Proestrus) vaginal impedance was additionally measured. Vaginal impedances greater than 3 kΩ were indicative of the proestrus stage. **B**, Schematic of hormonal fluctuations that are associated with the estrus-cycle (Neill, 2006). **C**, Fast-spiking neuron activity from a female out of estrus (in vaginal metestrus), line indicates period of social touch. **D**, Fast-spiking neuron activity from a female in estrus (vaginal estrus), line indicates period of social touch. **E**, Average ongoing firing rate relative to the day of vaginal estrus. Line represents the average value for each day, numbers represent the number of animals used to calculate the mean, shaded area represents the standard error of the mean. (One Way ANOVA, p=0.005; pairwise Tukey test: pre-estrus vs. estrus p=0.013, post-estrus vs. estrus p=0.011, pre-estrus vs. post estrus p=0.867). **F**, Average ongoing firing rate in and out of estrus (NE: N=20 animals, N=32 cells; E: N=9 animals, N=12 cells; p= 0.001, t-test). **G**, Regular-spiking neuron activity from a female out of estrus (vaginal diestrus), line indicates period of social touch, shaded area represents the standard error of the mean. **H**, Representative regular-spiking neurons from a female in estrus (vaginal estrus), line indicates period of social touch. **I**, Average ongoing firing rate of regular-spiking neurons relative to the day of estrus. Shaded area represents the standard error of the mean (Kruskal-Wallis p=0.694; pairwise Mann-Whitney *U* test: pre-estrus vs. estrus p=0.972, post-estrus vs. estrus 0.406, pre-estrus vs. post estrus p=0.594). **J**, Average ongoing firing rate of regular-spiking neurons in and out of estrus (NE: N=20 animals, N=31 cells; E: N=12 animals, N=25 cells; p= 0.626, Mann-Whitney *U* test). *p<0.05, **p<0.01, ***p<0.005, ****p<0.001.

The timing of the observed estrus-dependent modulation of fast-spiking neuronal activity corresponded to known timings of the increase in blood estrogen (Neill, 2006) (schematic **Figure 2B**). With this in mind, we hypothesized that estrogen may be responsible for modulation of fast-spiking neuronal activity. To test this idea, we obtained control of the female rat hormonal status by ovarectomizing females and externally supplementing estrogens. Specifically, we performed awake *in vivo* juxtacellular recordings from ovariectomized females that were fed either 17 beta-estradiol (28 μg/kg) or vehicle control (**Figure 3A**). Interestingly, we found that ongoing firing of fast-spiking neurons from ovariectomized females (OVX) was similar to firing in intact females out of estrus. When OVX females were fed estradiol, ongoing firing was significantly increased in single cells (**Figure 3B–C**) and on average across all fast-spiking neurons (**Figure 3D**). Much like the changes of ongoing activity in the estrus-cycle, the changes in ongoing firing of fast-spiking neurons were marked, with an approximately two-fold increase in firing rate. Responses to social touch also increased in single cells (**Figure 3B–C**) and the entire interneuron population when ovariectomized females were fed estrogen (**Figure 3E**). A responsiveness change was not seen in interneurons of freely cycling females (**Figure 2**). This difference could be attributed to unexpected variables introduced with ovariectomy or an incomplete reconstitution of female hormones. In contrast, ongoing firing and social touch responses did not change in regular-spiking neurons (**Figure 3F–I**). Together, these results show that somatosensory fast-spiking cortical neurons are potently and specifically modulated by the estrus-cycle and by estrogen.

**Figure 3:**
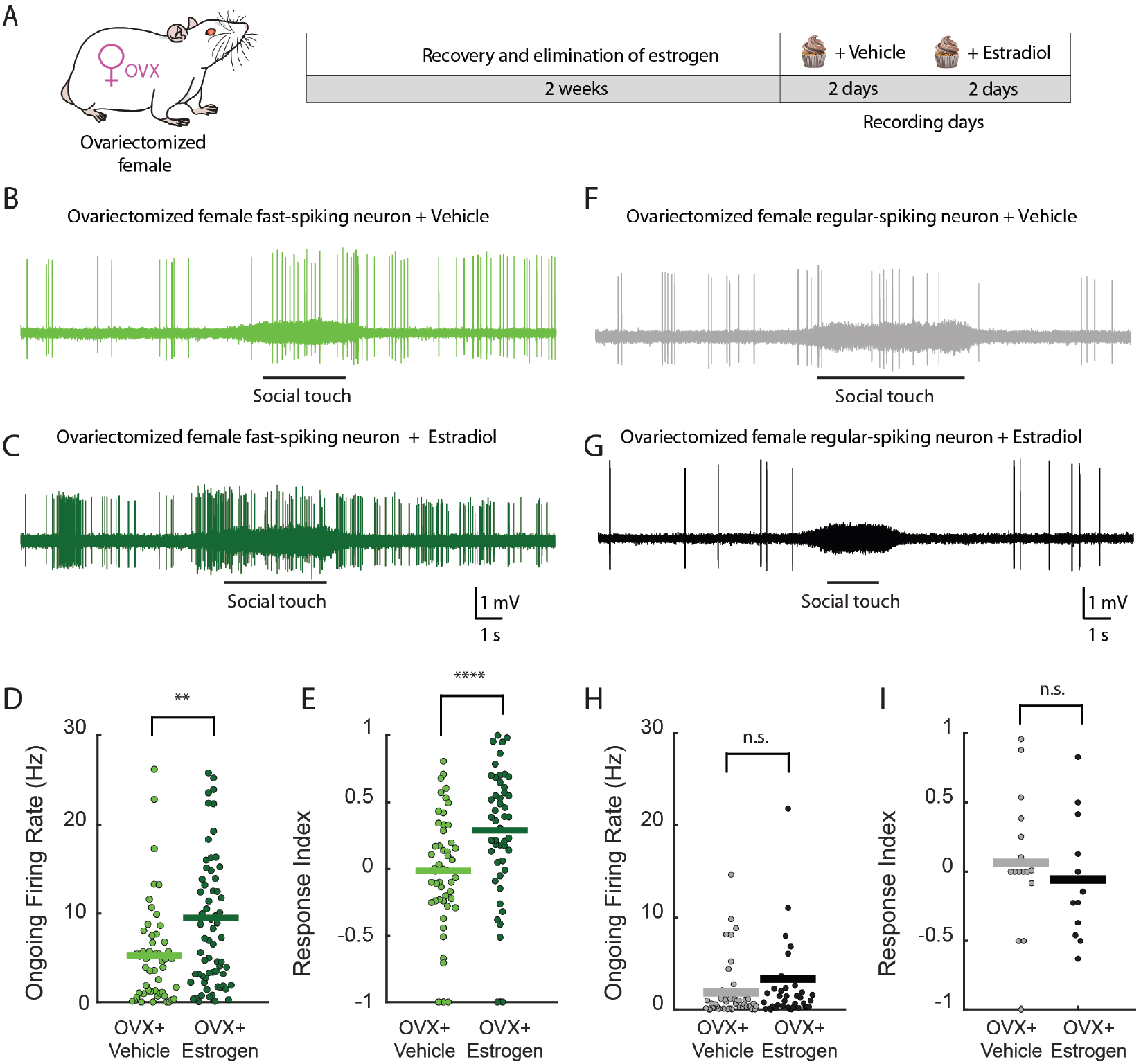
Estradiol increases fast-spiking neuron ongoing activity and responsiveness to social touch in ovariectomized females. **A**, Female rats were ovariectomized (OVX) prior to a 2 week waiting period to allow estrogen to be eliminated from the system. Subsequently, 4 days of juxtacellular recordings were performed (2 days where OVX females were fed nut cream with vehicle and 2 days where OVX females were fed nut cream with 17 beta-estradiol (E)). **B**, Fast-spiking neuron from an ovariectomized female fed vehicle. Gray area indicates time of social facial touch. **C**, Fast-spiking neuron from an ovariectomized female fed 17 beta-estradiol. Gray area indicates time of social facial touch. **D**, Ongoing firing rate of all recorded fast-spiking neurons from ovariectomized females fed vehicle or 17 beta-estradiol (OVX+Vehicle: N=51 cells; OVX+ Estrogen: N=67 cells; p=0.005 Mann-Whitney *U* test). **E**, Responsiveness all recorded fast-spiking neurons from ovariectomized females fed vehicle or 17 beta-estradiol (OVX+Vehicle: N=50 unique interactions OVX+Estrogen: N=49 unique interactions; p<0.001 Mann-Whitney *U* test). **F**, Regular-spiking neuron from an ovariectomized female fed vehicle. Gray area indicates time of social facial touch. **G**, Fast-spiking neuron from an ovariectomized female fed 17 beta-estradiol. Gray area indicates time of social facial touch. **H**, Firing rate of all recorded regular-spiking neurons from ovariectomized females fed vehicle or 17 beta-estradiol. (OVX+Vehicle: N=46 cells; OVX+Estrogen: N=39 cells; p=0.237 Mann-Whitney *U* test). **I**, Responsiveness of all recorded regular-spiking neurons from ovariectomized females fed vehicle or 17 beta-estradiol (OVX+Vehicle: N=16 unique interactions; OVX+Estrogen: N=13 unique interactions; p=0.487 Mann-Whitney *U* test). *p<0.05, **p<0.01, ***p<0.005, ****p<0.001.

A surprising aspect of our findings is that fast-spiking inhibitory neuron firing rates change without compensatory changes in regular-spiking excitatory firing rates in the female brain. This result is counter to a number of reports suggesting that excitation and inhibition in the cortex are balanced (Dehghani et al., 2016; Haider, 2006; Shu et al., 2003; Wehr and Zador, 2003). If fast-spiking inhibitory neurons change without comparable changes in excitation in females, what are the consequences and can such independent changes in excitation and inhibition be observed post-synaptically? To address these questions, we resolved excitatory and inhibitory postsynaptic potentials with *in vivo* whole-cell recordings from anesthetized adult female rats in and out of estrus. We recorded from putative and identified principal neurons while blocking spiking activity intracellularly (with inclusion of QX-314 2 mM in the internal solution) and depolarized neurons to reveal inhibitory events. We found an overall increase in the frequency of inhibitory post-synaptic potentials (IPSPs) (**Figure 4A– B, G**) with no change in IPSP amplitude (**Supplementary Figure 2A**) when females in estrus were compared with those out of estrus. In this case, the increase in IPSP frequency with estrus was also substantial (~ 1.4 fold), but smaller than the estrus-related changes we had observed in ongoing fast-spiking neuron activity. In this context, it is worth noting that fast-spiking neurons are only a fraction of cortical interneurons (Tremblay et al., 2016), hence, many of the of observed IPSPs presumably do not originate from fast-spiking neurons. Furthermore, it is also possible that the overall levels of ongoing inhibition may be reduced under anesthesia, which has been reported for other cortical areas (Haider et al., 2012). When excitatory post-synaptic events (EPSPs) were examined, no change in frequency (**Figure 4C–D, H**) or amplitude (**Supplementary Figure 2B**) was observed. To address the relative balance of inhibition to excitation in each cell, we then computed an I/E frequency index, where the frequency of inhibition was divided by the frequency of excitation and then normalized. This index further confirmed a shift in the balance of excitation and inhibition, where the contribution of inhibition was greater in estrus (**Figure 4I**). We next asked if responses to whisker stimuli measured with intracellular methods might also depend on the estrus state. We applied air puff stimuli to the whiskers of anaesthetized females and found that the latency to IPSP onset was significantly shorter in females in estrus (8.03 ± 0.69 ms, mean±sem) than the IPSP latency in females out of estrus (10.84 ± 1.08 ms, mean±sem, **Figure 4E–F**). The reduced inhibitory latency in estrus might result in a shortening of the integration time window, which could potentially lead to more precise coding during the estrus state. No change in evoked IPSP amplitude was observed (**Supplementary Figure 2C**), nor was a change in evoked EPSP latency or amplitude (**Supplementary Figure 2D–E**) observed. No change in input resistance or resting membrane potential was found across conditions (data not shown).

**Figure 4.**
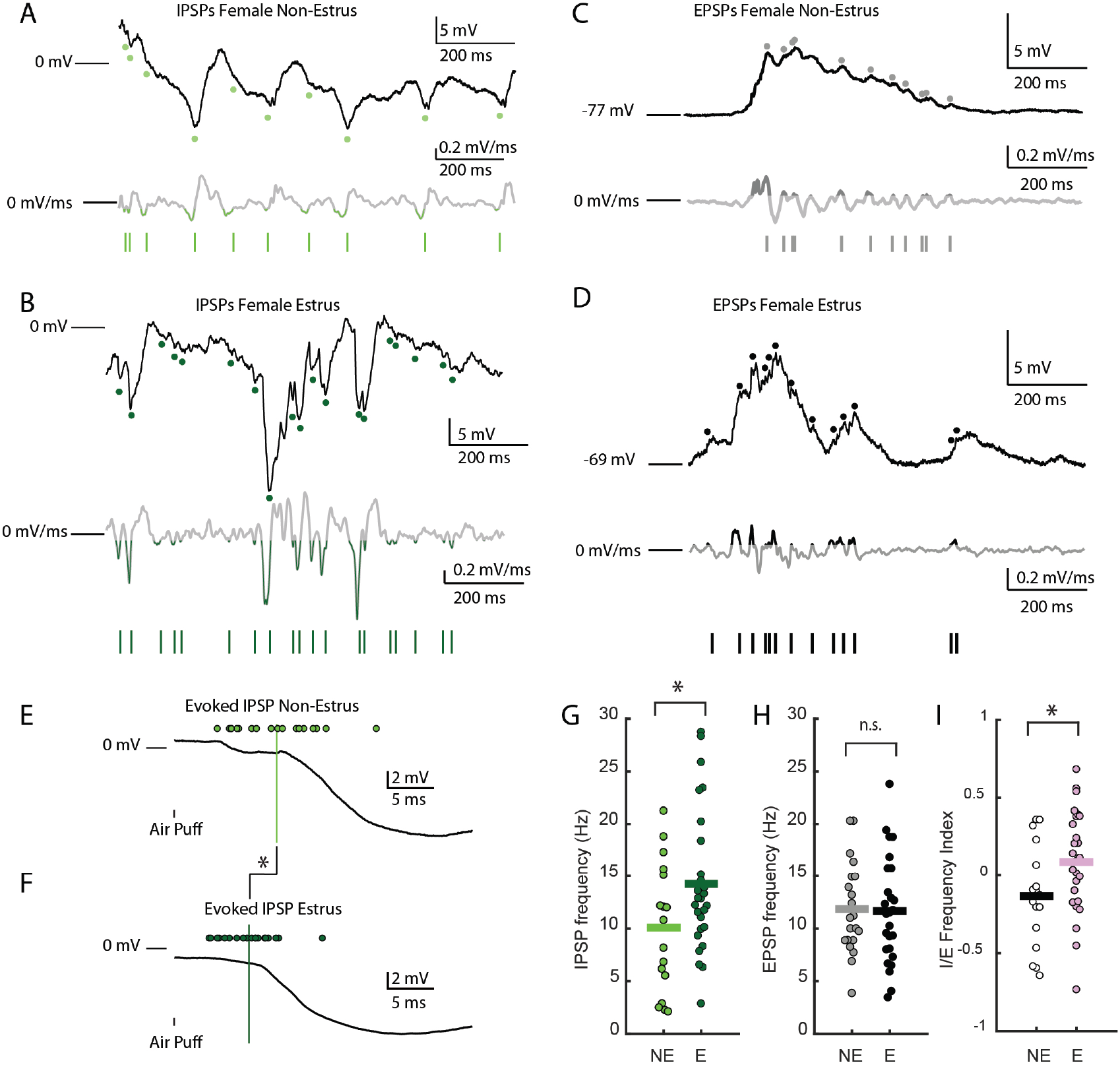
Estrus dependent changes in inhibition without changes in excitation. **A**, Inhibitory post-synaptic potentials (IPSPs) recorded from a putative principal neuron from a female out of estrus. QX-314 was included in the pipette to block spiking activity. The cell was depolarized with constant current injection to reveal post-synaptic inhibitory events (top). Events were detected as rapid changes in the voltage derivative (mV/ms, middle). Tick marks indicate IPSPs detected (bottom). **B**, IPSPs recorded from a female in estrus (top). Events were detected as rapid changes in the voltage derivative (mV/ms, middle). Tick marks indicate IPSPs detected (bottom). **C**, Excitatory post-synaptic potentials (EPSPs) recorded from a putative principal neuron from a female out of estrus. EPSPs were recorded at the cell’s resting potential (top). Events were detected as rapid changes in the voltage derivative (mV/ms, middle). Tick marks indicate EPSPs detected (bottom). **D**, EPSPs recorded from a female in estrus (top). Events were detected as rapid changes in the voltage derivative (mV/ms, middle). Tick marks indicate EPSPs detected (bottom). **E**, Evoked IPSPs recorded from females in (E: N=19 cells, green circles indicate latency of each cell) and **F**, out of estrus (NE: N=18 cells, green circles indicate latency of each cell; p=0.03, t-test). **G**, IPSP frequency for all cells in and out of estrus (NE: N=17 cells; E: 27 cells; p=0.046 t-test). **H,** EPSP frequency for all cells in and out of estrus (NE: N=21 cells; E: 26 cells; p=0.896 t-test). **I**, I/E frequency index (IPSP frequency/EPSP frequency) for all cells recorded in and out of estrus (NE: N=17 cells; E: 26 cells; p=0.038 t-test). *p<0.05, **p<0.01, ***p<0.005, ****p<0.001.

How does estradiol act to change cortical fast-spiking neuron activity? Does it act directly in the cortex or systemically (e.g. through extra-cortical projections)? We performed *in situ* hybridization for Esr2 messenger RNAs (estrogen receptor β) to assess co-localization with the fast-spiking neuron marker parvalbumin (PV) in the somatosensory cortex. We found that Esr2 mRNAs were found throughout cortical layers with a concentration in Layer 5. Highest co-localization of Esr2 mRNAs in PV neurons was also found in deep cortical layers (**Figure 5A–B**). Interestingly, when we ordered physiological recordings according to relative depth, we found that this mirrored relative levels of Esr2/PV co-localization (**Figure 5C**).

**Figure 5.**
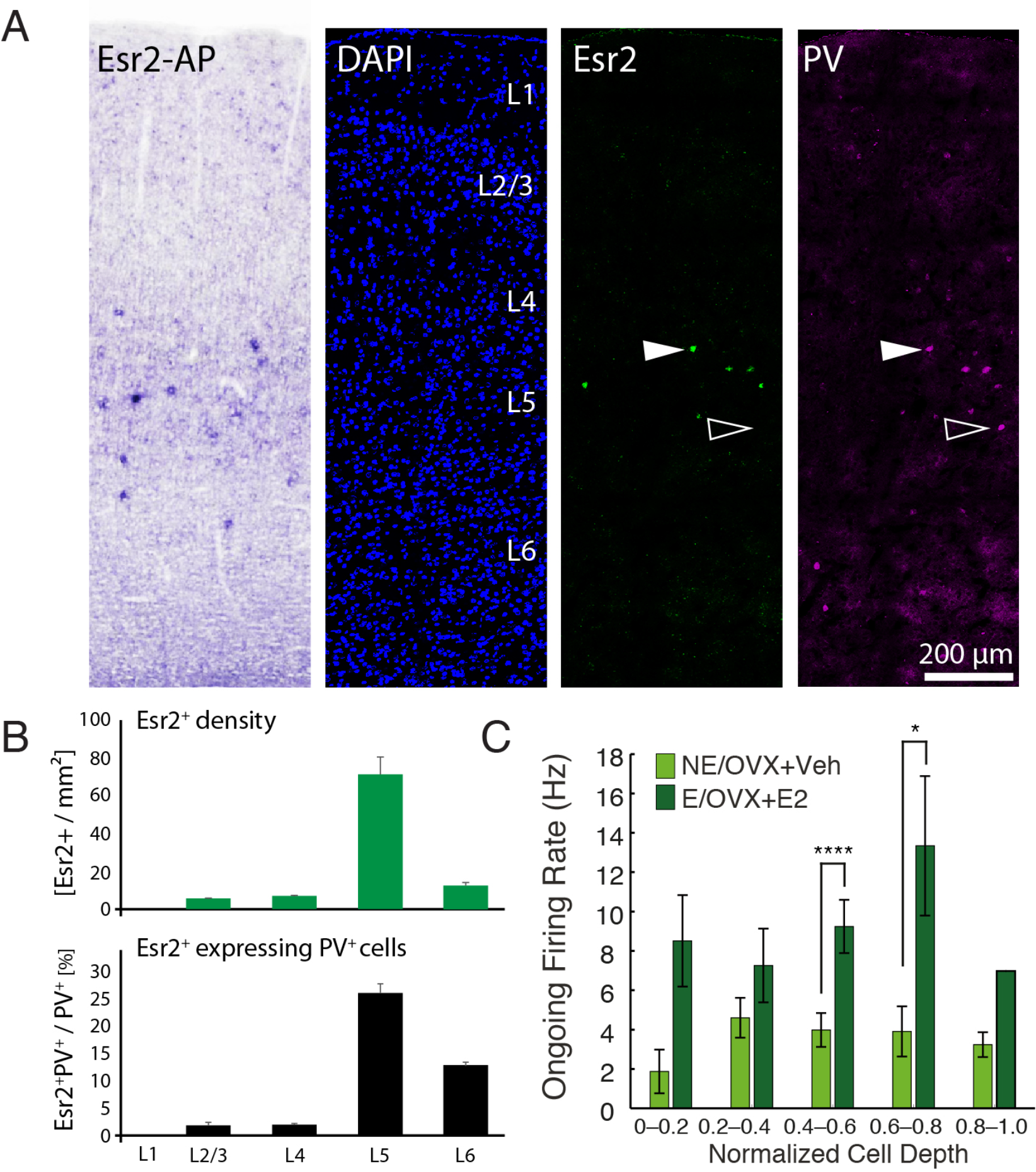
Co-localization of estrogen receptor β (Esr2) and parvalbumin (PV) in rat somatosensory cortex mirrors laminar distribution of fast-spiking ongoing firing rates in estrus and estrogen-supplemented conditions. **A**, (from left to right) In situ hybridization for Esr2 mRNA visualized with alkaline phosphatase staining shows that Esr2 is mainly expressed in layer 5 of somatosensory cortex of female rats. Layer 5 contains the highest number of highly expressing Esr2 cells, whereas weakly expressing cells may be also present in other layers. We only considered highly expressing cells for further analysis using fluorescent in situ hybridization (Esr2, green) combined with immunohistochemistry for parvalbumin (PV, purple). Examples indicated by white arrowheads are co-localization of Esr2 and PV, white open arrowheads are PV only cells. **B**, Highly expressing Esr2 cells are mainly located in layer 5 of somatosensory cortex, we find only few cells in other layers. The highest fraction of Esr2 and PV double positive cells is likewise located in layer 5, with some double positive cells also in layer 6. Graphs represent means ± S.E.M. **C,** Ongoing firing rates in Estrus and Estrogen supplemented states (where depth information was available) compared with Non-Estrus and Vehicle supplemented states (E/OVX+E2: 0–0.2 bin, N=6; 0.2–0.4 bin, N=19; 0.4–0.6, N=36; 0.6–0.8, N=12; 0.8–1.0, N=1; NE/OVX+Veh: 0–0.2 bin, N=3; 0.2–0.4 bin, N=37; 0.4–0.6, N=27; 0.6–0.8, N=8; 0.8–1.0, N=2; One Way ANOVA on Ranks p=0.004; E/OVX+E2 vs. NE/OVX+Veh: 0–0.2 bin, p=0.098 t-test; 0.2–0.4 bin, p=0.246 Mann- Whitney *U* test; 0.4–0.6, p<0.001 Mann-Whitney *U* test; 0.6–0.8, p=0.028 Mann- Whitney *U* test; 0.8–1.0, p=NA). Graphs represent means ± S.E.M. *p<0.05, **p<0.01, ***p<0.005, ****p<0.001.

We next applied 17 beta-estradiol to the cortex and performed *in vivo* juxtacellular recordings in anaesthetized ovariectomized females to address whether estrogen acts locally in cortical fast-spiking neurons (**Figure 6A**). We found a significant increase in ongoing firing rates when fast-spiking neurons were recorded after local estrogen application compared to vehicle application (**Figure 6B–C**; Mann-Whitney *U* test, p=0.013, N=9 Veh, N=14 E2, experiment and analysis performed blind to condition). To further elucidate the mechanism of estrogen action in the somatosensory cortex, we performed patch-clamp recordings in brain slices from ovariectomized female VGAT-Venus transgenic rats (Booker et al., 2014; Uematsu et al., 2008) (**Figure 6D**). Layer 5 VGAT-Venus cells were targeted with fluorescent illumination and characteristic fast-spiking firing patterns were confirmed prior to drug application. Slices underwent post-hoc processing to confirm that recorded cells were PV positive (**Figure 6E**). Cells were characterized in a 5–10 minute baseline period before 17 beta-estradiol (250 nM) was washed on and cells were subsequently characterized every 5 minutes for a period of 5–30 minutes. After Estradiol wash-in, cells showed a reduction of rheobase (**Figure 6F–G**) as well as an increase in firing frequency with 250 pA of current injection (**Supplementary Table 1**). No change in subthreshold membrane properties was observed (**Supplementary Table 1**), nor was a change in rheobase observed with vehicle wash-in (**Supplementary Table 1**). When experiments were performed in the presence of a specific estrogen receptor β antagonist (PHTPP, 1 μM), no change in rheobase was observed with 17 beta-estradiol wash-in (**Figure 6H–I, Supplementary Table 1**). Together, these results indicate that 17 beta-estradiol acts locally in deep layers of the cortex to increase excitability in fast-spiking parvalbumin-positive interneurons through an estrogen receptor β (Esr2) mechanism.

**Figure 6.**
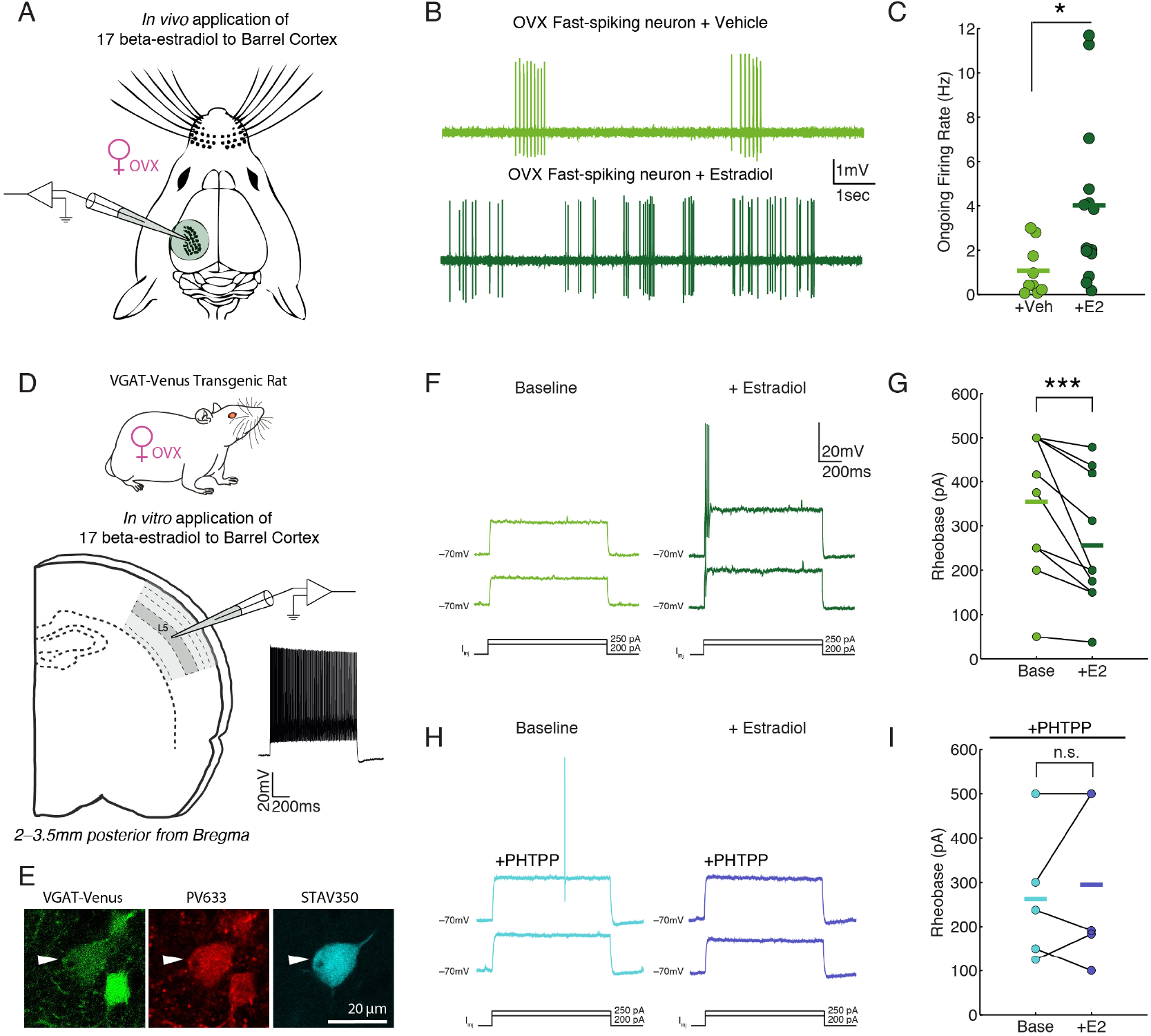
Action of 17 beta-estradiol occurs locally in the cortex through estrogen receptor β in parvalbumin-positive neurons. **A**, *In vivo* local application of 17 beta-estradiol to the surface of the cortex with juxtacellular recording in ovariectomized female rats. **B**, Example traces of fast-spiking neuron after vehicle application (above) and 17 beta-estradiol application (below) after 2 hours of diffusion. **C**, Ongoing firing rates of all recorded cells after vehicle or 17 beta-estradiol application to cortex (Mann-Whitney *U* Test, p=0.013, N=9 Veh, N=14 E2, experiment and analysis performed blind to condition). **D,** VGAT-Venus transgenic rats were ovariectomized 2 weeks prior to brain slice recording. Coronal brain slices were prepared 2–3.5 mm posterior from Bregma. Layer 5 cells were selected with fluorescent illumination and characteristic fast-spiking neuron firing patterns (example trace). **E**, Example of recorded cell which underwent post-hoc processing to confirm that it was parvalbumin-positive. VGAT-Venus signal (left), parvalbumin/Alexa 633 nm signal (middle), Streptdavidin 350 nm signal (right). **F**, Example of cell after baseline characterization (left) and after 250 nM 17 beta-estradiol wash-in (right). **G**, Change in rheobase after estradiol wash-in for all recorded fast-spiking cells (Wilcoxon Signed Rank Test, p=0.002 N=10). **H**, Change in rheobase after estradiol in the presence of estrogen receptor β blocker PHTPP (Passed Shapiro-Wilk normality test; Paired t-test, p=0.521, N=5). *p<0.05, **p<0.01, ***p<0.005, ****p<0.001.

## Discussion

Our data demonstrate that female cortical fast-spiking neurons are robustly activated with social touch, are cyclically regulated by the estrus state and are potently affected by administration of estradiol. We observed estrus-dependent changes in inhibition that were independent of changes in excitation as well as a reduction of evoked inhibitory latency *in vivo*. Using *in vivo* and *in vitro* electrophysiology recording methods and *in situ* hybridization, we further confirmed that the estrogen acts locally in the cortex to increase excitability in parvalbumin-positive neurons through an estrogen receptor β (Esr2) mechanism.

### Hormonal control and socio-sexual cortical information processing

Sex and sexual status have previously not played a major role in the analysis of information processing in primary sensory cortices. In recent years, however, evidence has accumulated that such variables have an impact on cortical responses. Analysis of social touch responses in the cortex of the rat revealed major differences in somatosensory cortical responses between male and female animals (Bobrov et al., 2014). Human studies have shown that differential responses in somatosensory cortex when social touch stimuli were initiated by male or female agents (although the actual social touch stimuli were identical) (Gazzola et al., 2012). Interestingly, it has been reported that the GABA_B_ agonist baclofen influences sexual receptivity in rats (Ågmo and Soria, 1997) although the contribution of the cortex to this finding is unknown. Evidence also points to an influence of ovarian hormones on cortical excitability in humans (Smith et al., 2002) and related observations to the ones made here come from the developing frontal cortex, where it was found that pre-pubertal (but not post-pubertal) hormones drive an increase in inhibitory neurotransmission (Piekarski et al., 2017).

### Cellular mechanisms

Our data (**Figure 3**) unequivocally show an impact of estrogens on cortical interneuron activity, which we have determined to occur through an estrogen receptor β (Esr2) mechanism in parvalbumin-positive interneurons (**Figure 5–6**). Immunostaining against estrogen receptor β and the calcium-binding protein parvalbumin has been previously reported (Blurton-Jones and Tuszynski, 2002; Kritzer, 2002), however this data has been challenged by the proven non-specificity of estrogen receptor β antibodies (Andersson et al., 2017; Snyder et al., 2010). We circumvented these concerns with the use of two *in situ* hybridization mRNA probes targeted to different regions of the Esr2 gene. Both probes labeled mRNAs in regions consistently identified as Esr2 positive including the paraventricular nucleus of the hypothalamus and medial amygdala (data not shown). Cortical expression was predominantly located in layer 5, however was consistently weaker than subcortical expression and required strong signal amplification, indicating low abundance of Esr2 mRNA. Therefore, we focused on highly expressing Esr2 cells and quantified their colocalization with parvalbumin and find the highest level of coexpression in layer 5 of somatosensory cortex. Although the estrogen receptor β (ERβ) is classified as a nuclear receptor, with traditionally postulated downstream effects on gene transcription, ERβ and the closely related ERα have been found in extra-nuclear sites in the brain including dendrites, spines, and axons (Milner et al., 2005; Woolley, 2007). Numerous studies have also reported that effects of estrogens in the brain can occur within the timeframe of minutes (Gu and Moss, 1998; Kumar and Foster, 2002; Wong and Moss, 1991; Woolley, 2007), which is suggestive of non-transcriptional effects. Our own *in vitro* slice data showed that 17 beta-estradiol initiates an increase in fast-spiking neuron excitability within minutes, which was then blocked with the selective ERβ receptor antagonist PHTPP (**Figure 6**). Together with our *in vivo* data in freely-cycling females, these findings support the notion that estrus-cycle driven estrogens have rapid and potent effects on fast-spiking neuronal excitability through a local effect on ERβ in parvalbumin-positive cortical neurons.

### The balance of excitation and inhibition

Despite the large number of studies reporting the balance (Dehghani et al., 2016; Shu et al., 2003; Wehr and Zador, 2003) and the precise timing (Okun and Lampl, 2008) of excitation and inhibition in the cortex, we are not aware of any that have closely examined possible sex differences or the influence of sex hormones in the adult cortex. Although earlier work has noted that excitation and inhibition are not balanced in all instances (Heiss et al., 2008; Isaacson and Scanziani, 2011) and that some E/I imbalances may influence social behavior (Yizhar et al., 2011) and enhance tuning for some sensory stimuli (Tan et al., 2007), the idea that carefully balanced excitation and inhibition generates stable cortical activity is still highly influential (Shadlen and Newsome, 1998). The present study, along with the work of other authors (Froemke, 2015; O’Donnell et al., 2017), suggest that the balance of excitation and inhibition is much more dynamic than previously thought and is unlikely to be the only pillar on which the stability of cortical activity rests. For example, negative feedback mechanisms such as the mutual inhibition of interneurons (Connors, 2017) might greatly contribute to the stability of cortical activity despite the absence of an accurate balance of excitation and inhibition. Although studies of cortical E/I balance in the female brain are few, it is interesting to note that a recent examination of estrogen-dependent development of inhibition was reported without a change in excitation (Piekarski et al., 2017). Additionally, changes in inhibition have been observed to occur independently of changes in excitation in other instances of neuromodulation such as by oxytocin (Owen et al., 2013) and noradrenalin (Martins and Froemke, 2015). Estrus related interneuron activity might act synergistically with changes in GABAergic receptor expression, as previously reported (Maguire et al., 2005), and these periodic changes in the tone of cortical inhibition might also contribute to the periodic occurrence of epilepsy in the menstrual cycle (Pennell, 2009). Furthermore, the observed shift in evoked inhibitory latency with estrus may have profound consequences for temporal integration and coding of incoming sensory stimuli (Pinto et al., 2000). Overall, these findings suggest that studies regarding the circuits of the cortex cannot be generalized across sexes. It is, thus, imperative that future examinations of cortical function regard the influence of sex and sex hormones.

## Acknowledgements

This work was supported by Humboldt-Universität zu Berlin, the BCCN Berlin, NeuroCure, the Gottfried Wilhelm Leibniz Prize (M.B.), a grant from the DFG (BR 3479/11-1, A.C. and M.B.), the Chinese Academy of Sciences President’s International Fellowship Initiative (Grant No. 2018PB0119, R.N.) and a grant from the NSFC (81741140, H.W.) and a grant from the NSFC (81741140, H.W.). We thank Viktor Bahr and Jean Simonnet for assistance with data analysis and Rajnish Rao for introduction to ovariectomy and vaginal smear procedures. We thank Christian Ebbesen, Johanna Sigl-Glockner, Jean Simonnet, Edith Chorev, Rick Gray and Rishi Narayanan for comments and discussion of the manuscript. We thank Dr. Imre Vida for generous contribution of VGAT-Venus transgenic female rats.

## Author contributions

A.C. and M.B. conceived the project. A.C. and C.L. collected and analyzed preliminary juxtacellular data together. A.C. performed and analyzed further awake and anaesthetized *in vivo* juxtacellular and whole-cell patch clamp data. P.B. and R.S. performed *in vitro* whole-cell patch clamp electrophysiology. A.C. and P.B. analyzed *in vitro* electrophysiology data. A.C. performed anatomical analysis, imaging and reconstruction of *in vivo* and *in vitro* recorded cells. A.C. and P.B. imaged *in vitro* recorded cells. L.L., H.W. and R.N. created probes, performed and analyzed *in situ* hybridization for Esr2. A.C. prepared the figures. A.C. and M.B. wrote the paper.

## Methods

### Animals

Experimental procedures were performed according to guidelines of the respective local ethics committees: German guidelines on animal welfare under the supervision of the local ethics committees (permit numbers: G0193/14, Brecht and T100/03, Schmitz) and the Animal Care and Use Committees at the Shenzhen Institute of Advanced Technology (SIAT), Chinese Academy of Sciences (CAS), China (permit number SIAT-IRB-171016-NS-WH-A0384).

### Assessment of Estrus Stage

The estrus stage of female rats (starting from P35) was determined by examination of vaginal smears (Marcondes et al., 2002) and with measurement of vaginal impedance (59160 Rat Vaginal Estrous Cycle Monitor, Stoelting Co., Wood Dale, IL). Both measurements were made at the same time daily, impedance measurements 3 kΩ and greater were considered to be the proestrus stage.

### Surgery and Implantation

#### Ovariectomy

Female Wistar and VGAT-Venus transgenic rats were ovariectomized at P35 under ketamine/xylazine anaesthesia (100 mg/kg ketamine and 7.5 mg/kg xylazine). Ovaries where removed from the dorsal side of the trunk following incision. Incision was sutured and rat was allowed to recover.

#### Chronic Awake Experiments

Ovariectomized and non-ovariectomized Wistar rats (P35–90) were handled for 2–3 days prior to surgical procedures. Animals were anesthetized with an initial injection of 100 mg/kg ketamine and 7.5 mg/kg xylazine. Respiration, blink and pinch reflex were observed throughout the surgery and, if needed, animals were injected with an extra shot (25%) of ketamine/xylazine mixture or a 25% dose of ketamine alone. Temperature was monitored using a rectal probe and maintained with a heating pad (Stoelting) set to 34–36°C. Lidocaine was locally injected in the scalp, which was then removed. During the first surgery a metal bolt was implanted on the skull of the animal’s head using an UV hardening glue (Kerr) and dental cement (Heraeus). After successful training of the implanted subject rat, a craniotomy was performed. The craniotomy was made by drilling a 1 to 1.5 mm diameter circle, which was centered 5.5 mm posterior and 2.5 mm lateral to bregma. The dura was left intact for awake juxtacellular recordings. After the craniotomy, the preparation was protected by implanting a threaded plastic cylinder with a removable lid. Between recording sessions, the preparation was covered using silicone (Kwik-Cast, World Precision Instruments).

#### Acute Anaesthetized Experiments

In acute electrophysiology experiments subject animals were anaesthetized with urethane (1.5-2 g/kg). One surgery was performed without prior habituation. Preparation of the craniotomy was identical to specifications for chronic experiments. In whole-cell patch clamp experiments and in juxtacellular experiments with local drug application, the dura was removed using a bent syringe and fine forceps.

### Habituation Procedure and Behavior for Awake Electrophysiology Experiments

After the first surgery, animals were allowed to recover overnight and a habituation procedure was performed, as previously described (Lenschow and Brecht, 2015). Two to three habituation sessions were performed per day starting with 5 minutes of head-fixation during the first session. Sessions were then increased by 10 minutes each day. The habituation period depended on the animal’s behavior but was typically 2–4 days. During habituation animals were gradually exposed to the experimental procedures and environment required for subsequent recordings. This included the addition of light, movement of the micromanipulators and the recording headstage. Stimulus rats were introduced to the subject rat in order to reduce stress at the time of experiments. Social interactions were staged by experimenters by presenting a stimulus rat to the headfixed subject rat (Figure 1A). Infrared lights were installed (Abus TV6830, wavelength 880 nm) above the recording setup with a 25 Hz low-speed (K240, Siemens) and 250Hz high-speed camera (Basler A504k). Behavior of the subject rat was monitored throughout the recording session using low-speed videography and the high-speed camera was triggered when social interactions took place. A social interaction (or “social touch”) was marked when the stimulus and subject rats touched whiskers. The beginning of an interaction was the first whisker overlap and the end of an interaction was the time-point at the end of whisker overlap. The timing of social touch episodes were scored offline using ELAN software (Max Planck Institute for Psycholinguistics, Nijmegen, The Netherlands) and later aligned to electrophysiological data using Matlab (MathWorks, Natick, MA).

### In Vivo Electrophysiology

Juxtacellular and whole-cell patch recordings were amplified (Dagan BVC-700A, Dagan, Minneapolis, MN), low-pass filtered at 10 kHz and sampled at 50 kHz by a data-acquisition interface (LIH 1600, HEKA or Power 1401, CED, Cambridge, England) controlled by Patchmaster (HEKA) or Spike2 software (CED, Cambridge, England).

#### Juxtacellular recordings

Patch pipettes containing intracellular recording solution were lowered into the cortex and cells were searched for using an audio monitor (Grass Technologies AM10) while steps were made in 4 μm increments with a micromanipulator (Luigs & Neumann SM-5). When spiking activity was detected, electrophysiological and video recordings of staged social interactions was performed (see above). Implanted females were recorded from daily while monitoring the estrus-cycle (up to 7 days) or as long as the preparation allowed further recordings.

#### Whole-cell recordings

Blind *in vivo* patch clamp recording (Margrie et al., 2002) were performed in females whose estrus-cycle was tracked for several days in advance to confirm cycle regularity and estrus stage was determined with vaginal smears and impedance measurement (see above). Patch pipettes were lowered into the cortex with positive pressure (200–300bar). When the pipette reached 150–200 μm below the surface, the pressure was lowered to 30 bars to search for cells. When the pipette resistance increased, suction was applied to establish a gigaohm seal and finally the whole-cell configuration. Whole-cell patch-clamp recordings were made by using glass electrodes made of borosilicate glass tubes (Hilgenberg). The resistance of the patch pipettes ranged from 4 to 6 MΩ. Pipettes were filled with an internal solution containing the following (in mM): K-gluconate 130, Na-gluconate 10, HEPES 10, phosphocreatine 10, MgATP 4, GTP 0.3, NaCl 4 and biocytin (~0.05%), pH 7.2. On the day of the recording, intracellular solution was thawed and 2 mM QX-314 (Tocris) was added to block voltage-gated Na^+^ conductances.

Upon establishment of a whole-cell recording, input resistance and resting membrane potential were noted. Several seconds of spontaneous activity were recorded at resting potential and when the cell was depolarized. For recordings of evoked EPSPs and IPSPs, air puff stimuli lasting 0.1 s were applied to the whiskers. Air puffs were generated from pulses of compressed air, delivered by a computer-triggered airflow controller (Sigmann Electronics, Germany). The air stimulus was delivered through a pipette which was positioned 3–4 cm from the whisker pad. The time from air puff stimulus to impact on the whiskers was estimated to be 28 ms, as determined by high-speed videography of a small piece of paper positioned at the whisker site.

### In Vitro Electrophysiology

#### Slice Preparation and Electrophysiology

Acute coronal slices of the somatosensory cortex from female VGAT-Venus rats (P47–87) were prepared at least 2 weeks after ovariectomy. Animals were anesthetized and decapitated. The brains were quickly removed and placed in ice-cold artificial cerebrospinal fluid (ACSF) (pH 7.4) containing (in mM) 87 NaCl, 26 NaHCO_3_, 25 glucose, 2.4 KCl, 7 MgCl_2_, 1.25 NaH_2_PO_4_, 0.5 CaCl_2_, and 75 sucrose. Tissue blocks containing the brain region of interest were mounted on a vibratome (Leica VT 1200, Leica Microsystems), cut at 300 μm thickness, and incubated at 35°C for 30 min. The slices were then transferred to ACSF containing (in mM) 119 NaCl, 26 NaHCO_3_, 10 glucose, 2.5 KCl, 2.5 CaCl_2_, 1.3 MgSO_4_, and 1.25 NaH_2_PO_4_.

The slices were stored at room temperature in a submerged chamber for 1–5 hr before being transferred to the recording chamber. The intracellular solution consisted of (in mM) 150 K-gluconate, 0.5 MgCl_2_, 1.1 EGTA, and 10 phosphocreatine (final solution pH 7.2).

#### Histology and Imaging of Labeled Neurons

On the last day of chronic *in vivo* experiments, in acute *in vivo* and *in vitro* slice experiments, neurobiotin (0.05–1%, Vector) was added to the internal pipette solution for post-hoc analysis of the recorded cells. The Pinault technique (Pinault, 1996) was applied to label and anatomically identify juxtacellularly recorded neurons. After successful *in vivo* recordings, animals were anaesthetized using a mix of ketamine and xylazine and perfused with phosphate buffer followed by a 4% paraformaldehyde solution (PFA). Brains were stored overnight in 4% PFA before preparing 80–150 μm coronal sections. For *in vitro* slice experiments, slices were fixed in 4% PFA overnight.

Neurobiotin-filled neurons were visualized using the avidin-biotin-peroxidase method (Vectastain ABC Kit, Vector) or using fluorescent streptdavidin (1:1000, Invitrogen streptavidin, Alexa 488, 350 or 546). After localizing labeled cells from *in vitro* and *in vivo* experiments, co-labeling was then performed with primary (1:1000, goat anti-Parvalbumin, Swant PVG-213) and secondary antibodies (1:500, Invitrogen Alexa secondary 546 or 633). Sections were visualized using a Leica epi-fluorescence microscope and an Olympus BX51 microscope (Olympus, Shinjuku Tokyo, Japan) equipped with a motorized stage (LUDL Electronics, Hawthorne NY) and a z-encoder (Heidenhain, Shaumburg IL, USA), was used for bright field microscopy and visualization of non-fluorescent (ABC processed) cells. Cells and processed were visualized and reconstructed using a MBF CX9000 (Optronics, Goleta CA) camera using Neurolucida or StereoInvestigator software (MBF Bioscience, Williston VT, USA). A Leica DM5500B epifluorescence microscope with a Leica DFC345 FX camera (Leica Microsystems, Mannheim, Germany) was used to image the immunofluorescent sections. Alexa fluorophores were excited using the appropriate filters (Alexa 350 – A4, Alexa 488 – L5, Alexa 546 – N3, Alexa 633 – Y5). Fluorescent images were acquired in monochrome, and color maps were applied to the images post acquisition. Co-labeled cells were additionally imaged with confocal microscopy (SP5 and SP8, Leica Microsystems, Mannheim, Germany) to ensure that there was no cross-talk between fluorophores.

### Analysis of Electrophysiological Data

#### Juxtacellular cell classification

Fast-spiking neurons (spike-width < 0.4 ms and Trough-to-Peak ratio > 0.3) and regular-spiking neurons (spike-width > 0.4 ms or Trough-to-Peak ratio < 0.3) were classified blind to condition. Classification criteria were established based on properties of morphologically identified neurons.

#### Analysis of in vivo whole-cell patch clamp data

Methods for analysis of spontaneous events were similar to those previously described (Okun and Lampl, 2008). Periods of spontaneous activity were filtered (Band-pass 3- 45 Hz) and the voltage derivative was computed at a time interval of 1 ms (25 points, sampling interval 0.04 sec). Subthreshold synaptic events were detected by finding rapid changes in the voltage derivative. The threshold for the voltage derivative was set just above the noise of each recording. If spikes were present, they were removed with spline interpolation. Spontaneous event amplitudes were computed by finding the baseline membrane potential 50 ms prior to the derivative threshold and subtracting it from the membrane potential value where the derivative threshold reversed. Analysis of spontaneous and evoked events was conducted blind to condition.

#### Analysis of in vitro whole-cell patch clamp data

Steady-state and transient input resistance, sag ratio, rheobase and firing rates were obtained from analysis of current clamp responses to a family of current injection steps, 1000 ms in duration, from –250 to 1000 pA. This characterization was repeated every 5 minutes during “Baseline” and post-Estradiol (“+E2”) periods (5–30 minutes after Estradiol wash-in). All sub- and supra-threshold properties were analyzed for each characterization and pooled across all baseline and post-Estradiol time points. Rheobase was classified as the current at which the first spiking occurred. Current steps where spontaneous spikes occurred and subsequent current injection did not elicit spikes were not classified as rheobase current steps. Steady-state and transient input resistance and the sag ratio were measured from voltage deflections in response to sub-threshold current steps. Steady-state input resistance was taken as the slope of voltage deflection (average of voltage in the last 500 ms of the trace) versus current injected. Transient input resistance was taken as the slope of voltage deflection (maximum initial voltage deflection) versus current injected. The sag ratio was calculated as the transient voltage deflection amplitude divided by the steady state voltage deflection amplitude. Firing rates (Hz) for each current injection step were taken from the number of detected spikes for each step (duration 1000 ms).

#### Software

Electrophysiological data was analyzed using Spike2 (CED, Cambridge, England), Igor Pro (WaveMetrics, Portland, OR) and custom written Matlab (MathWorks, Natick, MA) software.

### Pharmacology

#### In vivo experiments

Estrogen treated ovariectomized females were fed 17 beta-estradiol (28 μg/kg, Sigma Aldrich) which was dissolved in sesame oil and mixed with hazelnut cream. Vehicle treated ovariectomized females were fed sesame oil mixed with hazelnut cream alone. Both groups of animals were trained to eat the hazelnut cream alone, which they readily ate within 1–2 days. This method has been shown to restore physiological serum levels of 17 beta-estradiol in ovariectomized animals (Ström et al., 2012) and was advantageous to our awake recording scheme due to its non-invasive nature. In juxtacellularlocal application experiments, dura was removed and a chamber was implanted surrounding the craniotomy (see below). 17 beta-estradiol (100 μM, Sigma Aldrich) was dissolved in DMSO and then added to ringer solution (final DMSO concentration, 0.1%). 17 beta-estradiol or DMSO (“Vehicle”) was applied by filling the chamber and allowing 2 hours for diffusion to deep cortical layers.

#### In vitro experiments

17 beta-estradiol (250 nM, Sigma Aldrich) was dissolved in DMSO and added to external bath solution (final DMSO concentration, 0.01%). 17 beta-estradiol or DMSO (“Vehicle”) was washed on after baseline characterization of cells. The selective Estrogen Receptor Beta antagonist PHTPP (1 μM, Tocris) was dissolved in DMSO and diluted in external bath solution. Prior to recording, oxygenated slices were incubated in PHTPP for 30 minutes. Slices were then transferred to the recording chamber with PHTPP circulating for the course of the experiment. In the presence of PHTPP, 17 beta-estradiol (250 nM) was then washed on as before (see above).

### In Situ hybridization

#### Animals and Tissue preparation

Female adult SD rats (N = 3, 180-200 g; obtained from Beijing Vital River Laboratory Animal Technology Co., Ltd.) were used in the study. Animals were anaesthetized by isoflurane, and then euthanized by an intraperitoneal injection of an overdose of pentobarbital. Animals were then perfused transcardially with first 0.1 M phosphate buffered saline solution (PBS), followed by 4% formaldehyde, (from paraformaldehyde in 0.1 M phosphate buffer (PFA).

#### In situ hybridization and immunohistochemistry

In situ hybridization was performed as described before (Song et al., 2016). DIG-labeled Riboprobes were used for hybridization on 50 μm free floating cryosections. Hybridization was performed overnight at 65°C. Sections were washed at 65 °C twice in 2xSSC/50% Formamide/0.1% N-lauroylsarcosine, treated with 20 μg/ml RNAse A for 15 min at 37 °C, washed twice in 2xSSC/0.1% N-lauroylsarcosine at 37°C for 20 min and twice in 0.2xSSC/0.1% N-lauroylsarcosine at 37 °C for 20 min. Sections were blocked in MABT/10% goat serum/1% Blocking reagent (Roche, cat# 11096176001). For NBT/BCIP staining sections were incubated overnight with sheep anti-DIG-AP (1:1000, Roche cat# 11093274910). After washing, staining was performed using NBT/BCIP in NTMT until satisfactory intensity was reached. Fluorescent in situ hybridization was performed using TSA amplification. Sections were incubated with sheep anti DIG-POD (1:1000, Roche cat# 11207733910) and TSA reaction was performed using biotin-tyramide (ApexBIO). Subsequently sections were incubated with Streptavidin-Cy2 (Jackson ImmunoResearch) to detect the DIG labelled probe. Then we performed immunohistochemistry with antibodies against parvalbumin (1:2000, mouse anti-parvalbumin, Swant, V 235), which was detected with anti-mouse Alexa 594 (Jackson ImmunoResearch). Sections were washed and mounted with DAPI (Roti®-Mount FluoCare DAPI, Carl Roth).

Rat cDNA was synthetized from total brain RNA using EasyScript First-Strand cDNA Synthesis SuperMix (Transgen Biotech). We used PCR to obtain the DNA template employed for synthesizing the riboprobe from rat cDNA. We synthesized riboprobes for two parts of the rat Esr2 gene. Desired Esr2 fragments were amplified by PCR with the primer pairs indicated below (Phusion, NEB).

~~~
rEsr2-v1,   forward   1: agtaggaatggtcaagtgtggatccagg;   reverse   1: gagaaagaagcatcaggaggttggcc
rEsr2-v2, forward 2: atgaagtgtggtccctggacgc; reverse 2: aggagacaggtgcccagaagttgg.
~~~

PCR fragments were individually cloned in pEASY-Blunt Zero backbone (Transgen Biotech) and verified by sequencing. Antisense digoxigenin-labeled riboprobes were synthetized according to the protocol recommended by the manufacturer (Roche). Consistent with previous reports, both probes labelled cells in the paraventricular nucleus of the hypothalamus, medial amygdala, and several cortical areas including mainly layer 5 of the neocortex (Österlund et al., 1998; Shughrue et al., 1997). Subsequently, both probes were pooled together for staining and quantification.

#### Microscopy and counting

We acquired images using an inverted confocal microscope (LSM 880, Zeiss). Fluorescence images were acquired in monochrome and color maps were applied to images post-acquisition. Post hoc linear brightness and contrast adjustment were applied uniformly to the image under analysis. Esr2 fluorescence signal was quantified using ImageJ by measuring the Esr2 fluorescence in the DAPI positive region, with background subtraction. We classified cells into highly expressing and weak or non-expressing cells based on their levels of fluorescence intensity. Cells displaying signals higher than 1.5 fold of background levels were considered highly expressing and exclusively considered for this study. Cells with lower signal may express very low copy numbers of Esr2 mRNA, could not be reliably distinguished from background signals and were thus not considered for quantification. For neuron counting, 200 μm wide columns from the barrel cortex region were selected from coronal sections from three female rat brains. We then counted neurons manually using the CellCounter Plugin in ImageJ. We defined layers based on cell density in the DAPI channel and quantified Esr2, PV, and Esr2+PV neuron number on a per-layer basis.

### Statistics

For comparisons of two conditions, data were tested for normality (Shapiro-Wilk) and equal variance. When data were found to be normally distributed, a t-test (two-tailed) was performed. When data were non-normally distributed, a Mann-Whitney *U* test was performed. When groups of more than two were compared, a one-way analysis of variance (ANOVA) was performed for normally distributed data. For non-normally distributed data, a Kruskal-Wallis One Way ANOVA on ranks was performed.

Differences were considered statistically significant when p<0.05. Statistics were performed using SigmaPlot (Systat Software Inc., San Jose, CA) or in Matlab (MathWorks, Natick, MA).

### Data availability

The supporting data of all portions of the manuscript are available upon request from the corresponding authors.

## Supplementary Data

**Supplementary Figure 1.**
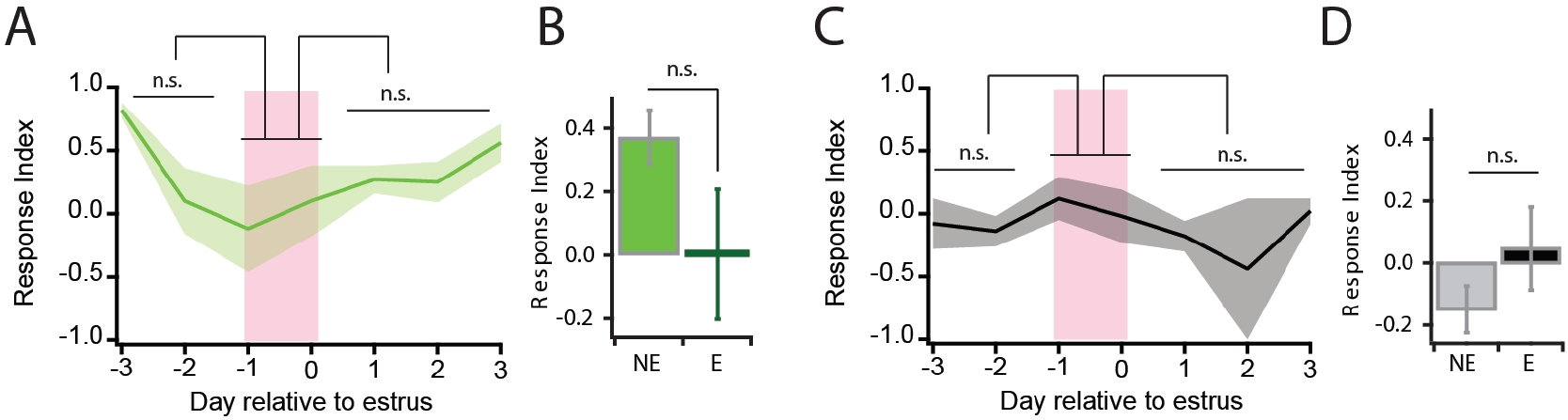
Juxtacellular responses to social touch in freely cycling fast-spiking and regular-spiking neurons. **A**, Average response index relative to the day of vaginal estrus in fast-spiking neurons. No significant change was observed (One Way ANOVA, p=0.161; pre-estrus vs. estrus: pairwise t-test p=0.187, post-estrus vs. estrus: Mann-Whitney *U* test p=0.229, pre-estrus vs. post estrus: p=0.765). **B**, Average response index in and out of estrus (NE: N=20 animals, N=32 cells; E: N=9 animals, N=12 cells; p=0.126 Mann-Whitney *U* test). **C**, Average response index relative to the day of vaginal estrus in regular-spiking neurons. No significant change was observed (One Way ANOVA, p=0.486; pre-estrus vs. estrus: pairwise t-test p=0.385, post-estrus vs. estrus: Mann-Whitney *U* test p=0.201, pre-estrus vs. post estrus: p=0.897). **D**, Average response index in and out of estrus in regular-spiking neurons (NE: N=21 animals, E: N=13 animals; p= 0.228, t-test).

**Supplementary Figure 2.**
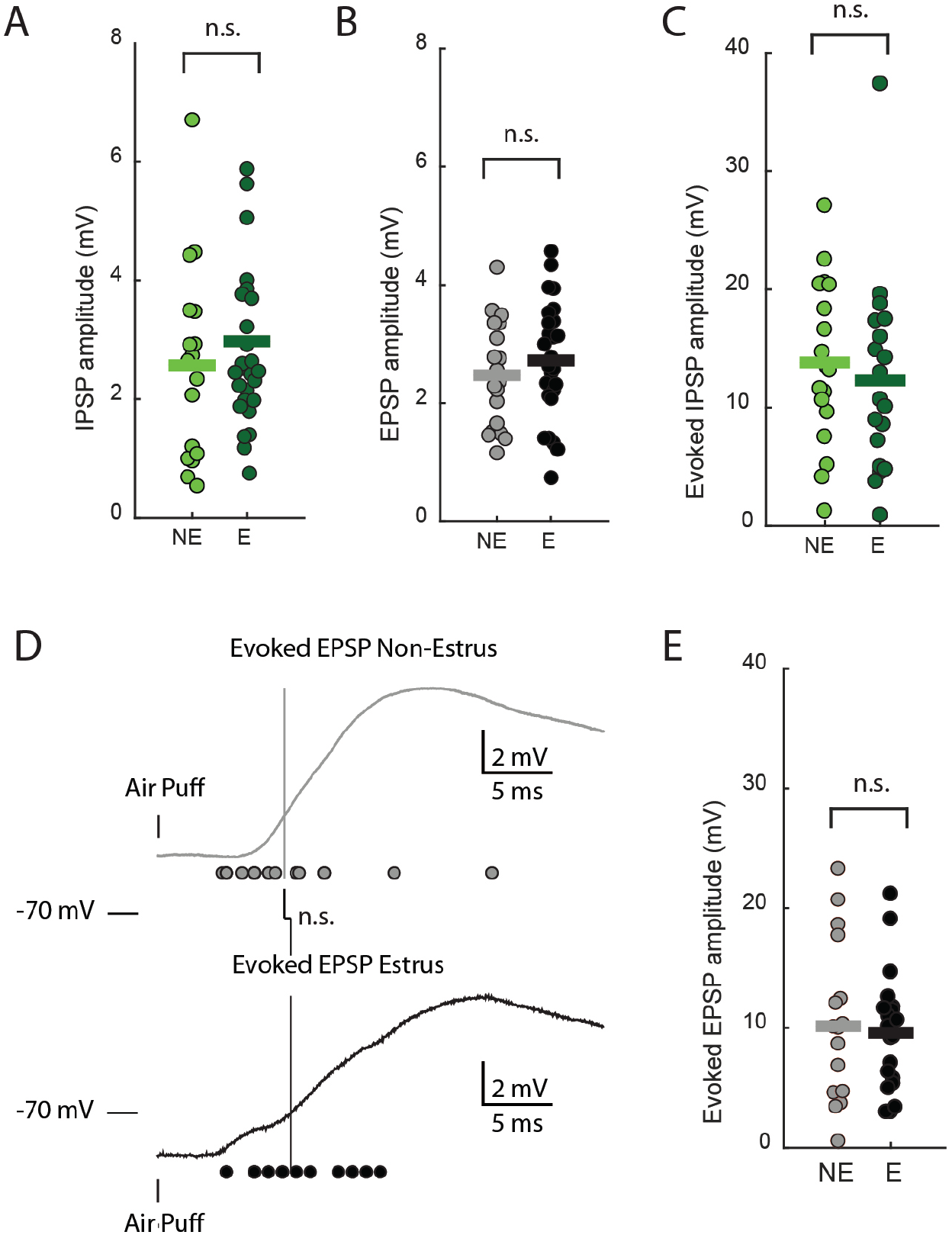
Supplemental intracellular figures. **A**, Spontaneous inhibitory post-synaptic potential amplitude (IPSP) is not different in and out of estrus. estrus (NE: N=17 cells; E: N=27 cells; p=0.539, Mann-Whitney *U* test). **B**, Spontaneous excitatory post-synaptic potential amplitude (EPSP) is not different in and out of estrus (NE: N=21 cells; E: N=26 cells; p=0.377, t-test). **C**, Evoked IPSP amplitude is not different in and out of estrus (NE: N=19 cells; E: N=22 cells; p=0.546, t-test). **D**, Evoked excitatory post-synaptic potential (EPSP) latency was not different in and out of estrus (NE: N=19 cells; E: N=22 cells; p=0.349, Mann-Whitney *U* test). **E**, Evoked excitatory post-synaptic potential (EPSP) amplitude was not different in and out of estrus (NE: N=19 cells; E: N=22 cells; p=0.779, Mann-Whitney *U* test).

**Supplementary Table 1:**
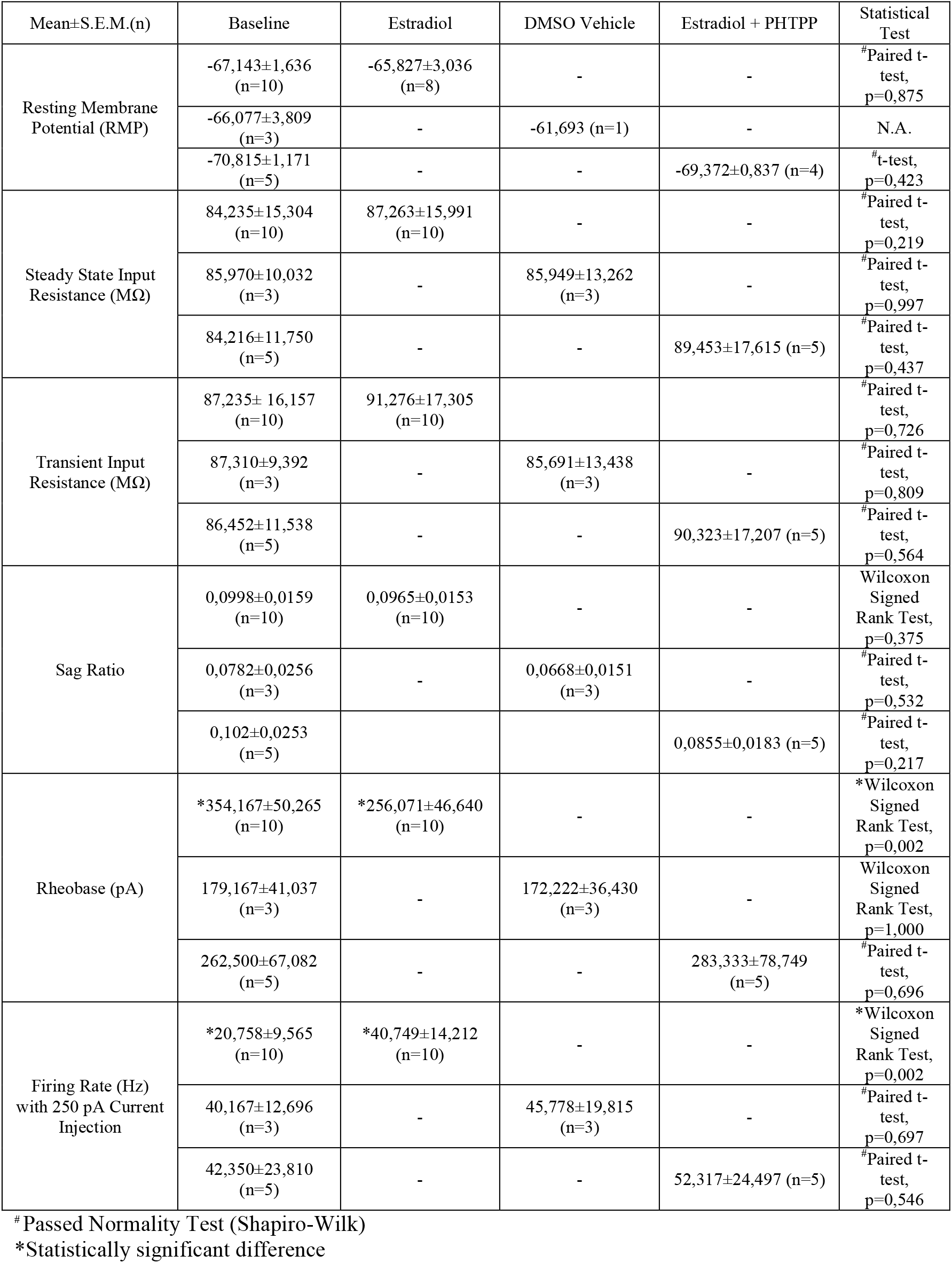
Analysis of *in vitro* electrophysiological data.

| Mean±S.E.M.(n) | Baseline | Estradiol | DMSO Vehicle | Estradiol + PHTPP | Statistical Test |
| --- | --- | --- | --- | --- | --- |
| Resting Membrane Potential (RMP) | -67,143±1,636 (n=10) | -65,827±3,036 (n=8) | - | - | <sup>#</sup>Paired t-test, p=0,875 |
|  | -66,077±3,809 (n=3) | - | -61,693 (n=1) | - | N.A. |
|  | -70,815±1,171 (n=5) | - | - | -69,372±0,837 (n=4) | <sup>#</sup>t-test, p=0,423 |
| Steady State Input Resistance (MΩ) | 84,235±15,304 (n=10) | 87,263±15,991 (n=10) | - | - | <sup>#</sup>Paired t-test, p=0,219 |
|  | 85,970±10,032 (n=3) | - | 85,949±13,262 (n=3) | - | <sup>#</sup>Paired t-test, p=0,997 |
|  | 84,216±11,750 (n=5) | - | - | 89,453±17,615 (n=5) | <sup>#</sup>Paired t-test, p=0,437 |
| Transient Input Resistance (MΩ) | 87,235± 16,157 (n=10) | 91,276±17,305 (n=10) | - | - | <sup>#</sup>Paired t-test, p=0,726 |
|  | 87,310±9,392 (n=3) | - | 85,691±13,438 (n=3) | - | <sup>#</sup>Paired t-test, p=0,809 |
|  | 86,452±11,538 (n=5) | - | - | 90,323±17,207 (n=5) | <sup>#</sup>Paired t-test, p=0,564 |
| Sag Ratio | 0,0998±0,0159 (n=10) | 0,0965±0,0153 (n=10) | - | - | Wilcoxon Signed Rank Test, p=0,375 |
|  | 0,0782±0,0256 (n=3) | - | 0,0668±0,0151 (n=3) | - | <sup>#</sup>Paired t-test, p=0,532 |
|  | 0,102±0,0253 (n=5) | - | - | 0,0855±0,0183 (n=5) | <sup>#</sup>Paired t-test, p=0,217 |
| Rheobase (pA) | *354,167±50,265 (n=10) | *256,071±46,640 (n=10) | - | - | *Wilcoxon Signed Rank Test, p=0,002 |
|  | 179,167±41,037 (n=3) | - | 172,222±36,430 (n=3) | - | Wilcoxon Signed Rank Test, p=1,000 |
|  | 262,500±67,082 (n=5) | - | - | 283,333±78,749 (n=5) | <sup>#</sup>Paired t-test, p=0,696 |
| Firing Rate (Hz) with 250 pA Current Injection | *20,758±9,565 (n=10) | *40,749±14,212 (n=10) | - | - | *Wilcoxon Signed Rank Test, p=0,002 |
|  | 40,167±12,696 (n=3) | - | 45,778±19,815 (n=3) | - | <sup>#</sup>Paired t-test, p=0,697 |
|  | 42,350±23,810 (n=5) | - | - | 52,317±24,497 (n=5) | <sup>#</sup>Paired t-test, p=0,546 |

<sup>#</sup> Passed Normality Test (Shapiro-Wilk)
\*Statistically significant difference

